# Expansion microscopy of nuclear structure and dynamics in neutrophils

**DOI:** 10.1101/2022.07.21.499684

**Authors:** Jason Scott Holsapple, Lena Schnitzler, Louisa Rusch, Tobias Horst Baldeweg, Elsa Neubert, Sebastian Kruss, Luise Erpenbeck

**Author notes:** Equal contribution (Jason Scott Holsapple, Lena Schnitzler).

## Abstract

Neutrophils are key players of the immune system and possess an arsenal of effector functions, including the ability to form and expel neutrophil extracellular traps (NETs) in a process termed NETosis. During NETosis, the nuclear DNA/chromatin expands until it fills the whole cell and is released into the extracellular space. NETs are composed of DNA decorated with histones, proteins or peptides and NETosis is implicated in many diseases. Resolving the structure and dynamics of the nucleus in great detail is essential to understand the underlying processes but so far super-resolution methods have not been applied. Here, we developed an expansion microscopy-based method and determined the spatial distribution of chromatin/DNA, histone H1, and nucleophosmin (NPM1) with a 4.9-fold improved resolution (< 40 nm) and increased information content. It allowed us to identify the punctate localization of NPM1 in the nucleus and histone-rich domains in NETotic cells with a size of 54 nm. The technique could also be applied to components of the nuclear envelope (lamins B1 and B2) and myeloperoxidase (MPO) providing a complete picture of nuclear dynamics and structure. In conclusion, expansion microscopy enables super-resolved imaging of the highly dynamic structure of nuclei in immune cells.

**Why it matters:** Accessibility to high-resolution imaging is critical to advancing research across various disciplines. However, conventionally this requires demanding optical hardware, special fluorophores or data analysis. Expansion microscopy is a technique adaptable to different cell and tissue types and is comparatively inexpensive and easy to perform. Applying this technique to cells and compartments such as the nucleus of immune cells that are difficult to image due to their size and morphology, yields valuable structural insights that would otherwise require more difficult super-resolution methods.

## Introduction

Neutrophilic granulocytes are an essential part of the innate immune system and comprise the most abundant type of granulocytes, making up 40% to 70% of all white blood cells in humans. In many ways, neutrophils possess exceptional properties that allow them to migrate quickly to the place of inflammation, squeeze through the endothelium and initiate immune responses^1–3^.

Over the last years, it has become increasingly clear that the rate-limiting factor for cellular mobility is nuclear morphology and the biomechanics of nuclear deformation. Neutrophils possess an exceptionally deformable nucleus with a unique composition of the nuclear envelope. Mature neutrophils lack lamins A/C, which have been shown to be essential for mechanotransduction during confinement^4,5^, and only contain low amounts of lamin B1 and B2^6–8^, which may constitute the mechanistic basis for the morphological plasticity of the neutrophil nucleus. Furthermore, the exceptional composition of the neutrophil nucleus also facilitates the formation of NETs^9–11^, an immune defense mechanism during which the entire neutrophil chromatin decondenses and expands first within the nuclear envelope and then, after rupture of said envelope, within the cytoplasm^12^. Ultimately, the neutrophil chromatin, decorated with a multitude of antimicrobial peptides and enzymes, is expelled into the extracellular space, where it can immobilize and eliminate diverse pathogens such as bacteria, fungi or even viruses^13^. When NET formation becomes dysregulated, however, it is implicated in several conditions including cancer metastasis, autoimmune diseases^14–16^, and even severe COVID-19^17^. There are many factors that can modulate this process, including adhesion and substrate elasticity^9^, ultraviolet light exposure^10^, and the presence of serum proteins such as albumin^11^. The physical properties of chromatin also importantly contribute to the formation and release of the NET due to the entropic swelling that occurs during the decondensation phase of NETosis^7,13,14^.

Despite the apparent importance of the nuclear composition for its immune defense functions and for the formation of NETs, studying nuclear functions and morphology remains difficult, as this requires very high-resolution imaging up to super-resolution, which is technically challenging, time-consuming and costly. Of note, neutrophils are comparatively small cells (approx. 10 μm diameter) with an even smaller and anisotropic nucleus (2 μm). Thus, changes in the distribution of chromatin and nuclear proteins during NETosis are yet to be fully understood.

Super-resolution methods have seen a tremendous progress in the past years and include STED^19–21^, PALM, STORM^22^, SOFI^23^ and Miniflux^24^. These methods enable resolving biological structures far below the optical resolution limit but are complex in terms of the optical hardware or the necessary data processing^25–31^. An alternative method is to increase the distance between fluorophores by expanding the sample (expansion microscopy). Expansion microscopy is a novel method wherein cells or tissues are embedded in a matrix and isotropically expanded^32–35^. This process produces a nearly transparent sample with a gain in resolution that renders conventional microscopy a viable means to obtain super-resolved images^36–39^. This has been used to explore intracellular dynamics but also to further understand the interface between cells and their attachment matrix^40^.

Here, we have developed a method to analyze neutrophils and their organelles such as the nucleus by expansion microscopy, achieving up to around 5-fold higher resolution than the Abbe limit. The nucleus of neutrophils is especially interesting for such methods because, as outlined above, it is highly heterogeneous and dynamic. We used this approach to study the distribution of histone H1 as well as lamins B1 and B2 within the nuclear envelope, nucleophosmin (NPM1) and to image characteristic neutrophilic molecules such as MPO. Furthermore, we characterized neutrophil chromatin composition and histone distribution in unstimulated cells and in neutrophils undergoing NETosis. Along with a higher resolution, expansion microscopy of neutrophils yields a higher spatial separation of nuclear structures, revealing more details about their distribution and dynamics.

## Materials and methods

### Neutrophil isolation

All experiments with human neutrophils were approved by the Ethics Committee of the University Medical Center (UMG) Göttingen (protocol number: 29/1/17). Neutrophils were isolated from fresh venous blood of healthy donors. Beforehand, all donors were fully informed about possible risks and the informed consent obtained in writing, consent could be withdrawn at any time during the study. Blood was received in S-Monovettes EDTA (7.5 ml, Sarstedt) and neutrophils isolated according to previously published standard protocols^12,41^. Neutrophils were resuspended in 1 mL HBSS^-Ca2+/Mg2+^. Cells were counted and further diluted at the required concentration for the following procedures in RPMI 1640 containing 10mM HEPES (Roth) and 0.5% human serum albumin (Sigma Aldrich). Purity of the isolation was assessed by a cytospin assay (Cytospin 2 Zentrifuge, Shanson) and Diff Quick staining (Medion, Diagnostics). Cell purity was always greater than 98%.

### NET induction

Fresh isolated human neutrophils were seeded in two 8-well, glass bottom chamber slides with removable chamber (ThermoFisher Nunc Lab-Tek II Chamber Slide System) at 80,000 cells/well in 200 μl RPMI 1640 (Lonza) containing 10mM HEPES (Roth) and 0.5% FCS (Biochrom GmbH, Merck Millipore). One chamber slide was used for gel expansion and the other one as non-expanded control. The chamber slides were incubated for attachment for 30 minutes at 37 °C. For NET formation, cells were activated with 100 nM phorbol 12-myristate 13-acetate (PMA, Sigma Aldrich) for a defined period (15 minutes, 30 minutes, 60 minutes, as indicated) whilst incubated at 37 °C and 5% CO_2_. To stop NETosis, cells were fixed with 3.2% paraformaldehyde (PFA) and 0.1% glutaraldehyde in phosphate buffered saline (PBS, Sigma Aldrich) as final concentrations. The fixed samples were washed with PBS 200 μl/well. Then, cells were incubated in 100 mM glycine solution for 5 minutes at room temperature. Cells were washed again twice with PBS and stored over night at 4 °C.

### Immunofluorescence staining

Before the staining procedure, the silicone sealing gasket was removed from the chamber slides. To block unspecific antibody binding, cells were incubated with blocking/permeabilization buffer (PBS with 0.1% Triton X-100 and 15% BSA) for 30 minutes at room temperature. Subsequently, cells were stained with polyclonal antihuman lamin B1 (IgG, rabbit, 1:50) (ab16048, Abcam), monoclonal anti-human lamin B2 (IgG2a, mouse, 1:100) (MA5-17274, Invitrogen), polyclonal anti-human MPO (IgG, sheep, 1:50) (AA16-718, Antibody online), monoclonal anti-human Histone H1 (IgG2a, mouse, 1:100) (MA5-13750, Invitrogen), or monoclonal anti-human NPM1 (IgG1, mouse, 1:500) (325200, Invitrogen) in blocking/permeabilization buffer for 120 minutes, washed three times with PBS and visualized with polyclonal anti-rabbit Alexa 488 (IgG, goat, 1:50) (ab11034, Abcam), polyclonal anti-mouse Alexa 488 (IgG, goat, 1:2000) (A11029, Invitrogen), polyclonal anti-mouse Alexa 555 (IgG, goat, 1:50) (A21422, Life Technologies), or anti-sheep Alexa 568 (IgG, donkey, 1:50) (ab175712, Abcam) as secondary antibody in blocking/permeabilization buffer for 60 minutes at 37 °C. After three more washes with PBS, chromatin was stained with Hoechst 33342 (1:2000 in PBS) (Thermo Fisher) for 15 minutes at room temperature. The nonexpanded probes were mounted with Faramount Mounting Medium (Dako Agilent Technologies) on a cover slip. The probes for expansion were mounted with PBS and stored over night at 4 °C.

### Expansion microscopy

The protocol was modified from the previously published protocol from Chozinski et al. 2016^42^. Neutrophils were prepared as described above. The silicone sealing gasket of the chamber slide was removed from the microscope slide. Blocking and staining was carried out as described above. Stained cells were fixed with 0.25% glutaraldehyde in PBS for 10 minutes at room temperature. After washing in PBS, cells were incubated in monomer solution (1x PBS, 2M NaCl, 2.5%/0.15% acrylamide/N’N’-methylenebisacrylamide, 8.625% sodium acrylate) for one minute at room temperature. Gelation chamber was assembled by attaching three coverslips with water drops on a glass slide, forming a three-sided chamber. Gelation solution was prepared by adding 10% TEMED and 10% APS to the monomer solution to final concentrations of 0.2%. Next, a drop of 1000 μL gelation solution was quickly put into the chamber. The chamber slide was laid slowly onto the drop, the cell-side facing the drop and the edges resting on top of the chamber-coverslips. The solution was allowed to gelate for 30 minutes at room temperature. Subsequently, the gel was removed together with the chamber slide from the gelation chamber, cut into 8 pieces, corresponding to the former 8 wells. Each gel was incubated in digestion buffer (8 U/mL proteinase K in 1x TAE (Tris base, acetic acid, EDTA), 10% Triton X-100, 8% guanidine HCl) for 30 minutes at 37 °C. The gels were then expanded in distilled water. The water was exchanged 4 times in 30-minute intervals. After the last exchange, the gel was placed on a 35 mm imaging glass bottom dish. The samples were imaged with the Axiovert 200 or the Axio Imager M1 microscope or with a confocal laser scanning microscope (Olympus IX83 inverted microscope, software: Olympus Fluoview v.4.2; see below). Area and eccentricity of the nuclei was measured with ImageJ. Eccentricity was calculated as:

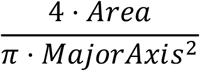

### Confocal microscopy

For confocal images an Olympus IX83 inverted microscope (software: Olympus Fluoview Ver.4.2, Olympus, objective UPLFLN60XOI) was used. Hoechst fluorescence was excited at 405 nm, lamin B1, histone H1, and NPM1 fluorescence at 488 nm, and lamin B2 and MPO fluorescence at 568 nm. All pictures were further processed with ImageJ (version 1.53g, National Institutes of Health) and MATLAB (version R2020a, The MathWorks, Inc.).

### Parameter calculations

All image processing for parameter calculations was conducted with Python. First, single cells from the microscope images were cropped, then a threshold was applied to the DNA-stained images to create a mask which only included staining intensities of the neutrophils and blocked all noise. These masks were applied to the corresponding DNA and Histone images. These images could then be used for parameter calculation. We also calculated the binning of the expanded microscope images by averaging 4×4 pixels to one, to simulate the effect of the at least 4 times lower resolution of nonexpanded microscope images in comparison to expansion microscopy images. The Pearson coefficient was calculated with the Python SciPy library (scipy.stats).

### Statistics

The colocalization of DNA and Histone was calculated with the Pearson’s coefficient. For all statistical significance tests, either the Mann-Whitney U test (using the scipy.stats package in Python, version 3.8.5, 64-bit) or unpaired t-test (using GraphPad Prism 5, version 5.04, 95% confidence intervals) was performed. The letter “n” indicates the number of independent experiments from individual donors. For every donor and all conditions, at least 40 cells were evaluated in a blinded manner.

## Results

### Expansion microscopy of primary human neutrophils and NETs

Building upon previous protocols for expansion microscopy of cultured cells and histological samples^43,44^, we developed a technique for the staining and visualization of single primary human immune cells and specifically for neutrophilic granulocytes.

**Figure 1:**
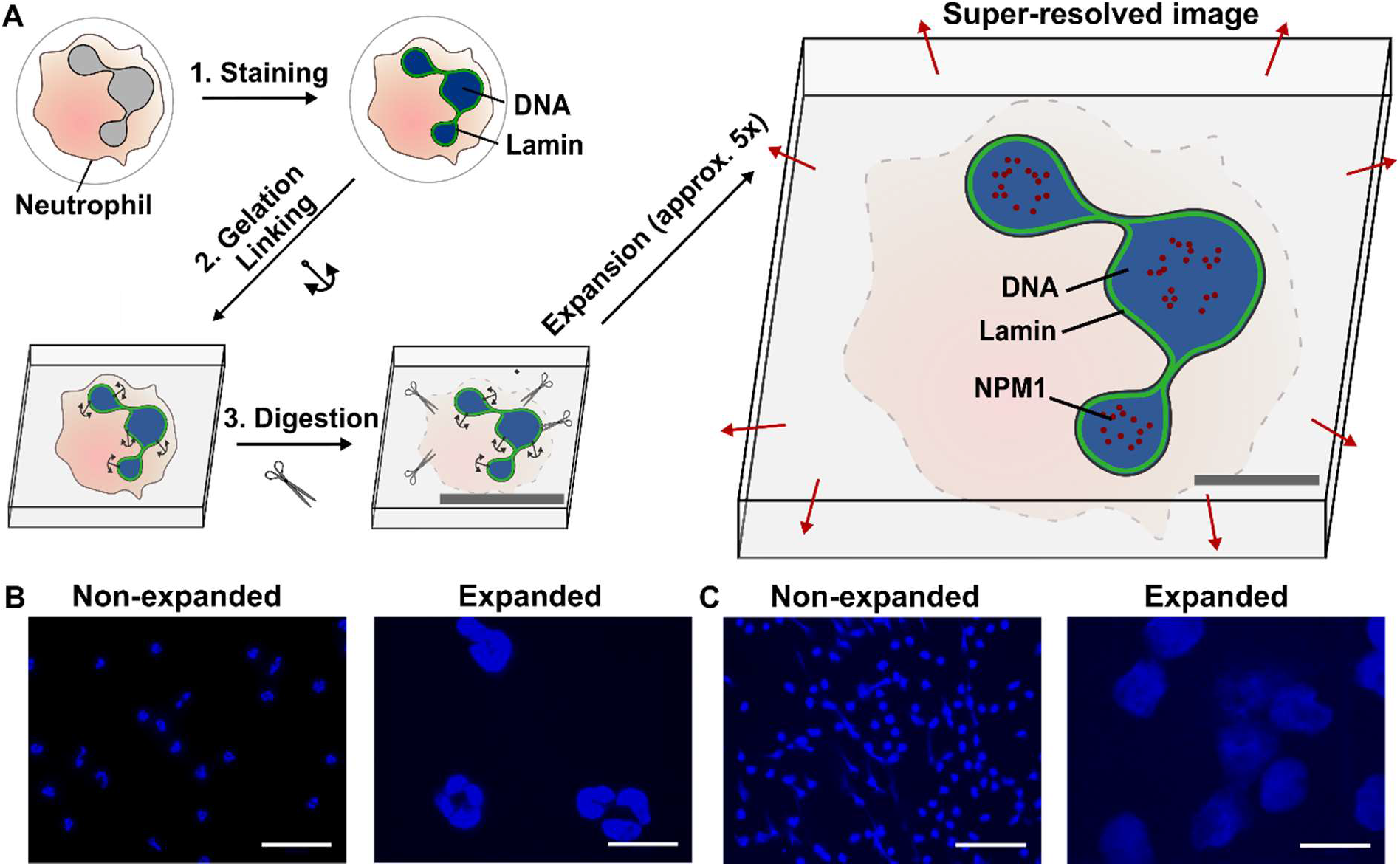
Expansion Microscopy of neutrophils and NETs. Schematic of the method (**A**): Cells are stained with fluorescently labeled antibodies or common fluorescence dyes (e.g., Hoechst). They are then embedded in a polyacrylamide gel and proteins are digested. Afterwards the gel is isotropically expanded via swelling in distilled water. Representative confocal microscopy images of Hoechst-stained nonexpanded or expanded neutrophils (**B**) and NETs (**C**). Scale bars = 10 μm (**A**), 50 μm (**B**), 100 μm (**C**).

Freshly isolated human neutrophils were placed in chamber slides and activated to form NETs. They were then fixed and stained by using standard immunofluorescence techniques. Following a second fixation step with glutaraldehyde, cells were then engulfed in monomer solution and the gelation process initiated (Figure 1A) on top of a glass coverslip. The resulting gel samples were incubated in digestion buffer, containing proteinases, and finally placed in distilled water to initiate the swelling process. After overnight expansion of the gel, cells within the sample were ready to be visualized either by conventional widefield or confocal microscopy (Figures 1B and 1C). Alternatively, by using 8-well chamber slides for the initial settling of the neutrophils and then later cutting the gel into 8 pieces according to the previous borders of the chambers, it was also possible to perform up to 8 different stainings of the cells in the respective chambers.

After establishing the general feasibility of this protocol, we assessed the magnification we were able to reach, using freshly isolated neutrophils that were stained with the nuclear dye Hoechst. In one dimension, a 4.9 fold magnification was achieved for the stained nuclei, amounting to a 25-fold increase in chromatin-stained area in the 2D images (Figure 2A). To study cellular morphology with this novel method, it is important that the proportions are maintained during mechanical enlargement of the previously stained structures. For this reason, we also assessed whether expansion of the nuclei was isotropic, and thus that it is uniform in all directions. To this end, we determined eccentricity of the stained nuclei before and after expansion and found that eccentricity did not vary between expanded and non-expanded nuclei (Figure 2B). We thus confirmed that proportions of stained structures were not altered by expansion microscopy.

In addition to neutrophil nuclei, we also stained NETs by Hoechst and observed them after following the protocol of expansion microscopy. As NETs are composed of strands of decondensed chromatin decorated with a variety of peptides and proteins, they are very fragile and often subject to mechanical artifacts during staining procedures. Here, we could show that NETs could be visualized similarly well as nonexpanded nuclei using the above-explained staining, fixation and expansion procedure. A 4- to 5-fold magnification was again achieved by expansion (Figure 2A), while eccentricity of NETs did not change after the procedure (Figure 2B). Thus, expansion microscopy is suited even for the visualization of fragile cellular structures.

**Figure 2:**
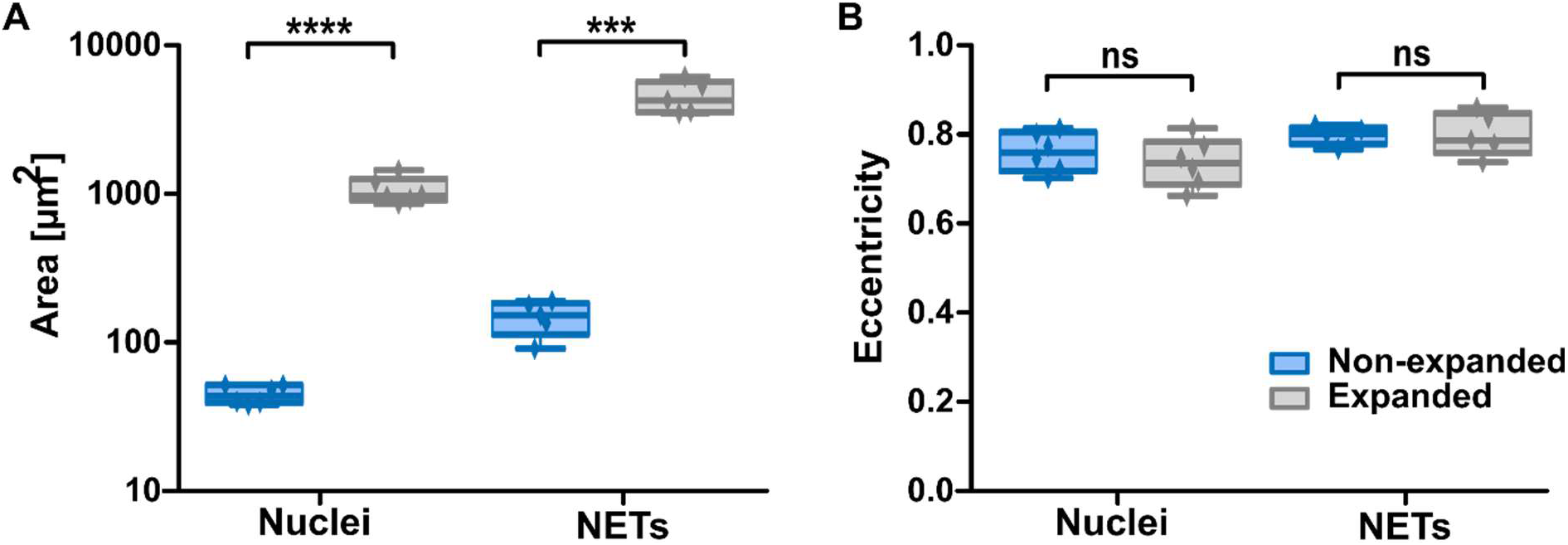
Isotropy of expansion of neutrophil nuclei and NETs. With expansion, the areas of neutrophil nuclei (n = 6) or NETs (n = 5) increases (**A**) in contrast to eccentricity of nuclei or NETs, which is constant (**B**). The mean nuclear area increases from 44.8 μm^2^ to 1060 μm^2^, amounting to a lateral expansion of 4.9. Boxplot shows interquartile range, mean values and minimum to maximum (whiskers). Significance was tested using unpaired t-tests, *** = p≤0.001, **** = p≤0.0001.

### Expansion microscopy significantly improves resolution and provides more information on spatial distribution of nuclear structures

Next, we assessed whether expansion microscopy provided measurable improvements over non-expanded cells. To this end, we analyzed line scans through representative images of unstimulated neutrophils that had been stained for DNA, histone H1, and NPM1, and subsequently processed in Python (Figure 3). The line scan through neutrophil nuclear lobules revealed intensity signals for all three stainings throughout the cell. In unstimulated neutrophil nuclei, chromatin staining was moderately enhanced towards the periphery of the nucleus, while histone H1 localized strongly towards the periphery of the nucleus.

In contrast, NPM1 was distributed throughout the lobules of the nucleus. As a theoretical construct to assess the advantages of expansion microscopy, we calculated binned images of the expanded microscope images by averaging an area of 4 by 4 pixels to one, to simulate the effect of the at least 4-times lower resolution of nonexpanded microscope images in comparison to expansion microscopy images.

**Figure 3:**
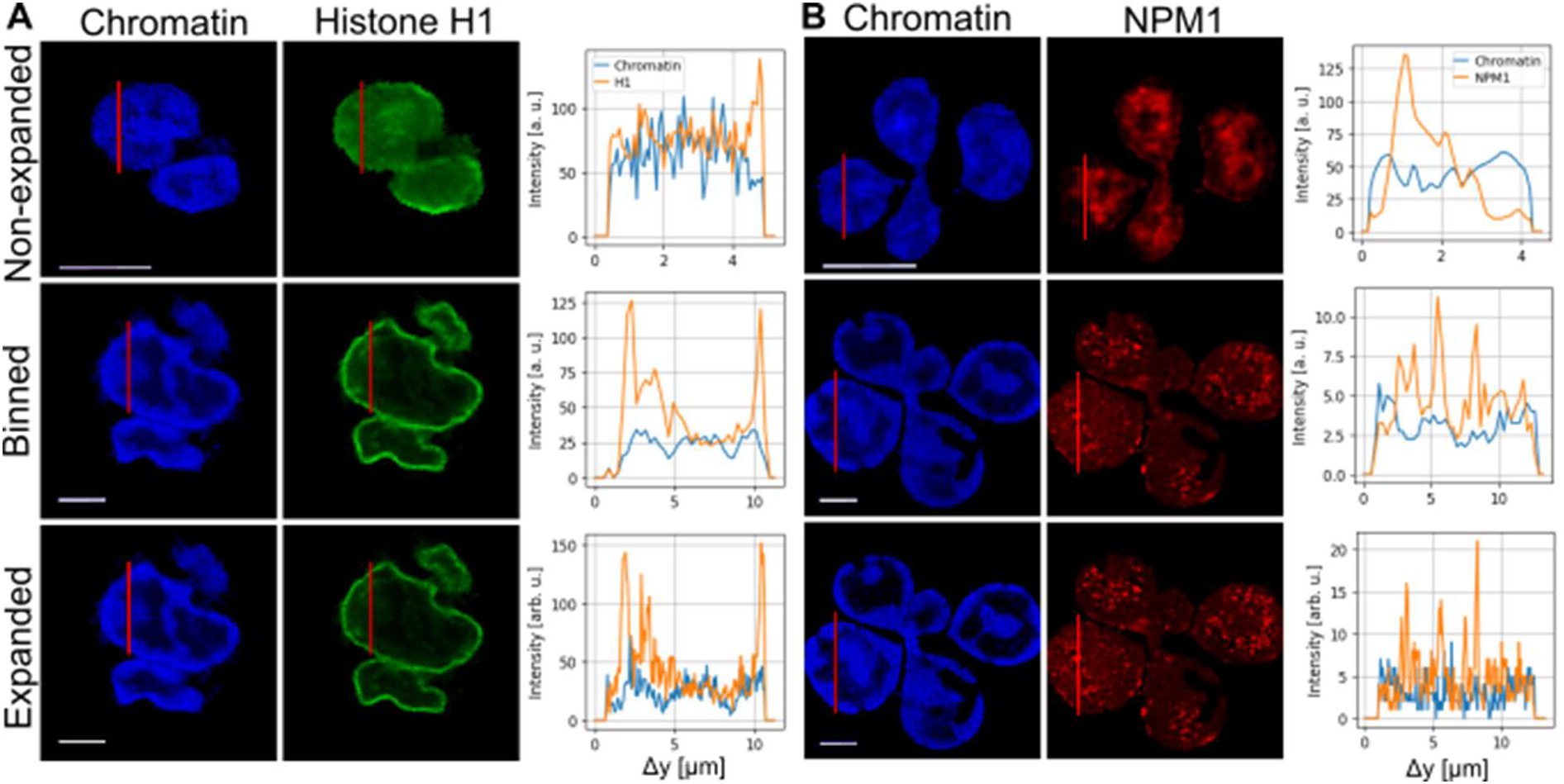
Distribution of chromatin and nuclear features in expanded and non-expanded neutrophils. (**A**) Fluorescence images of Hoechst (left) and histone H1 (middle) labelled neutrophils without expansion microscopy, binned (4×4) expansion microscopy images, and with expansion microscopy. The red line indicates a line scan through the nucleus and is shown on the right. Note that the cells shown for nonexpanded are not the same as for expanded/binned. All images show unstimulated cells. (**B**) Fluorescence images of Hoechst (left) and NPM1 (middle) labelled neutrophils without expansion (top) microscopy, binned (4×4) expansion microscopy images (middle), and with expansion microscopy (bottom). The results indicate that there is more spatial information contained in the higher-resolution expansion microscopy images. All scale bars are 5 μm for expanded and non-expanded.

In the line-scans (Figure 3), an increase of resolution by expansion microscopy images compared to normal, non-expanded microscopy and to the binned cells is clearly visible by the higher density of information and the increase in details on intensity distribution. It is important to note that the expansion factor cannot be determined by comparing the length of the line scans of different cells due to the heterogeneity of the individual lobes. This would only be applicable in comparing the same fixed cell pre- and post-expansion.

In a next step, colocalization of DNA and histones was then calculated using the Pearson’s coefficient from the statistical functions of the Python SciPy library (Figure 4). Colocalization was assessed in unstimulated expanded, non-expanded and “binned” cells (see above). A lower coefficient indicates a higher degree of spatial separation between stained structures. Of note, in the expanded images a significantly lower Pearson’s coefficient was shown for expanded versus non-expanded cells, reflecting the above-described enhancement of histone H1 at the border of the chromatin-stained area (as shown in Figures 3A and 5). In conclusion, non-expanded images provided much less information about distinct localization of DNA and histones, as can be derived from the Pearson’s coefficient that was close to 1. For the binned images, Pearson’s coefficient was in the same range as for the non-expanded cells, as was to be expected because information regarding colocalization is lost in the binning process. Since the Pearson’s coefficient of binned and non-expanded images are in the same range, the artificial loss induced with the binning seems also to be similar to the 4-fold difference in resolution between expanded and non-expanded images.

**Figure 4:**
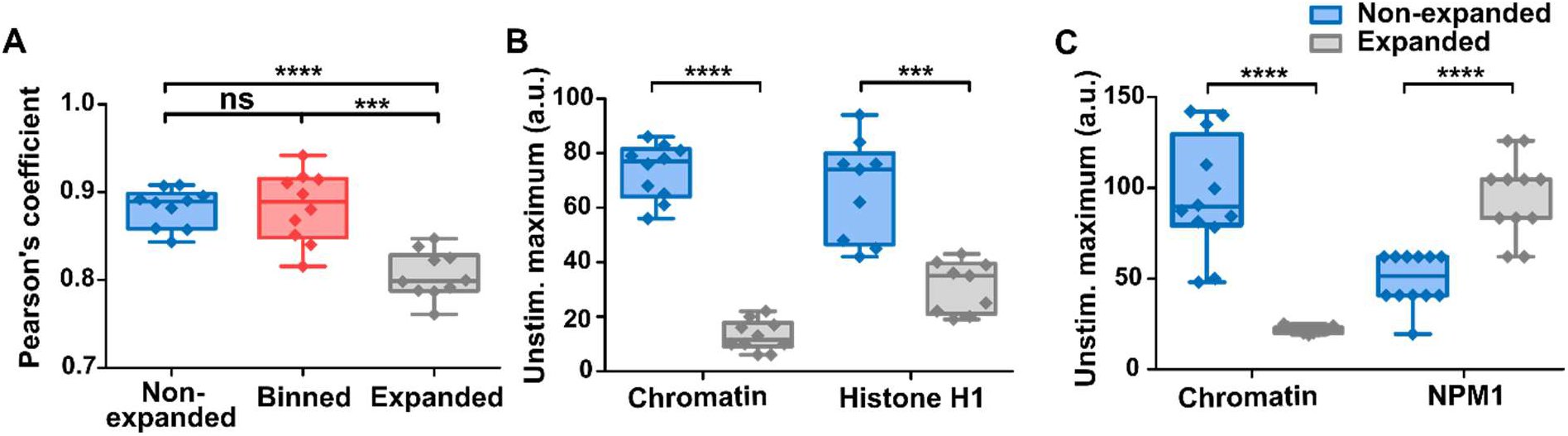
Impact of expansion on information content and maximum signal intensity. (**A**) The Pearson’s coefficient serves as a measure for colocalization for unstimulated cells stained with Hoechst and H1 for expanded, binned (4 pixels) and non-expanded images. Colocalization is smaller for expanded images indicating more information content. Statistical analysis was performed with the Mann-Whitney U test. p-values: *** = 0.001, **** = 0.0001, n.s. = not significant (n_expanded ≥ 10, n_non-expanded ≥ 10). (**B**, **C**) Maximum signal intensity decreases with expansion in unstimulated cells stained for chromatin and histone H1 (**B**), while these values are relatively lower overall for NPM1. (**C**) Statistics were performed with the Mann-Whitney U test. p-values: *** = 0.001, **** = 0.0001, n.s. = not significant (n_expanded ≥ 9, n_non-expanded ≥ 9). Boxplot shows interquartile range, mean values and minimum to maximum (whiskers).

Expansion dilutes the density of fluorophores, which could decrease the contrast of the image. In case of the Hoechst dye the mean fluorescence intensity (at the same imaging conditions) decreased (Figure 4B) like the expansion factor (4.9). In contrast, the antibody-based histone H1 staining decreased much less, indicating a certain level of proximity-based quenching of the fluorophores. In case of NPM1 the fluorescence signal of the expanded sample is even higher, which means that the loss of proximity-based quenching outcompetes dilution because of expansion. This result can be explained by the punctuate high-density localization of NPM1.

### Increase of spatial information in biological processes like NETosis

Next, we studied NETosis by conventional microscopy and by expansion microscopy, after staining for DNA (Hoechst) and histone H1 (Figure 5). The images revealed a preferential distribution of histone H1 towards the periphery of the nucleus, particularly at the interface between chromatin and the cytoplasm. Chromatin was similarly enriched towards the periphery of the nucleus, although less prominently compared to H1. During the process of NETosis, chromatin starts to expand within the cell^45^. Typically, this also entails mixing of nuclear and cytoplasmic proteins with the chromatin. Interestingly, the histone H1 staining in expanded cells revealed the emergence of additional “interfaces” with an enhancement of H1 staining at 60 min (Figure 5). In non-expanded images, this was not visible, illustrating the gain of biological information by expansion microscopy.

In cells that are completely filled with DNA just before membrane rupture (Figure 5, first row on the right), histones and DNA showed a rather even distribution of chromatin and histone H1. This agrees with previously performed STED microscopy that showed no fine structure of the chromatin^12^. However, the higher resolution achieved with expansion enabled us to reveal a certain granularity of the histone H1 and DNA. These domains had a diameter 266 nm ± 45 nm (expanded), which corresponds to 54.3 nm ± 9.2 nm before expansion (Supplementary Figure 3). It suggests that the DNA meshwork is not homogenous but rather like a soft material with incorporated more rigid spheres (‘raisins in a cake’) and this should have implications for the mechanical properties of the cells.

**Figure 5:**
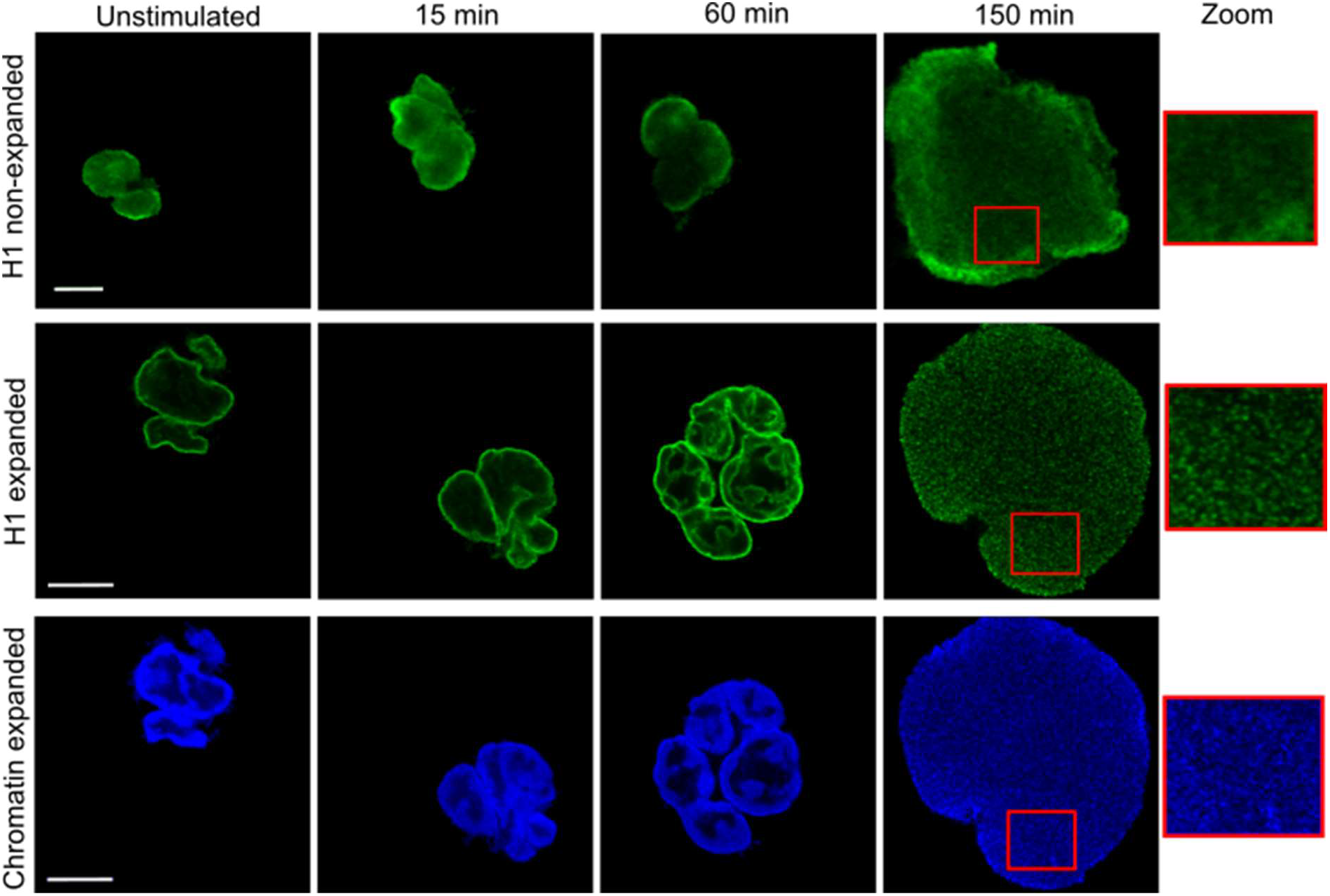
Identification of histone-rich domains below the resolution limit. Exemplary images of neutrophils undergoing NET formation after activation with 100 nM PMA. DNA was stained by Hoechst (bottom row) and histone H1 (upper and middle rows). In the first row, non-expanded cells are shown as a comparison to the expanded cells in the second and third rows. DNA and histone H1 colocalize during the first part of NET formation. For NETotic cells before membrane rupture (150 minutes) an enlarged ROI is shown, illustrating granularity and histone-rich domains with higher DNA density. Scale bars 10 μm for expanded and 5 μm for non-expanded images.

### Expansion microscopy is suited for the visualization of diverse cytoplasmic and nuclear neutrophil structure

To further assess the suitability of expansion microscopy of neutrophils for different cellular structures, we performed immunofluorescence stainings for cytoplasmic proteins (MPO) and components of the nuclear lamina (lamin B1 and B2). The staining of histone H1 and DNA had already been performed for Figures 3 and 5. We were able to perform all immunofluorescence stainings and increase the resolution for both cytoplasmic as well as nuclear markers as seen in Figures 6 and 7.

**Figure 6:**
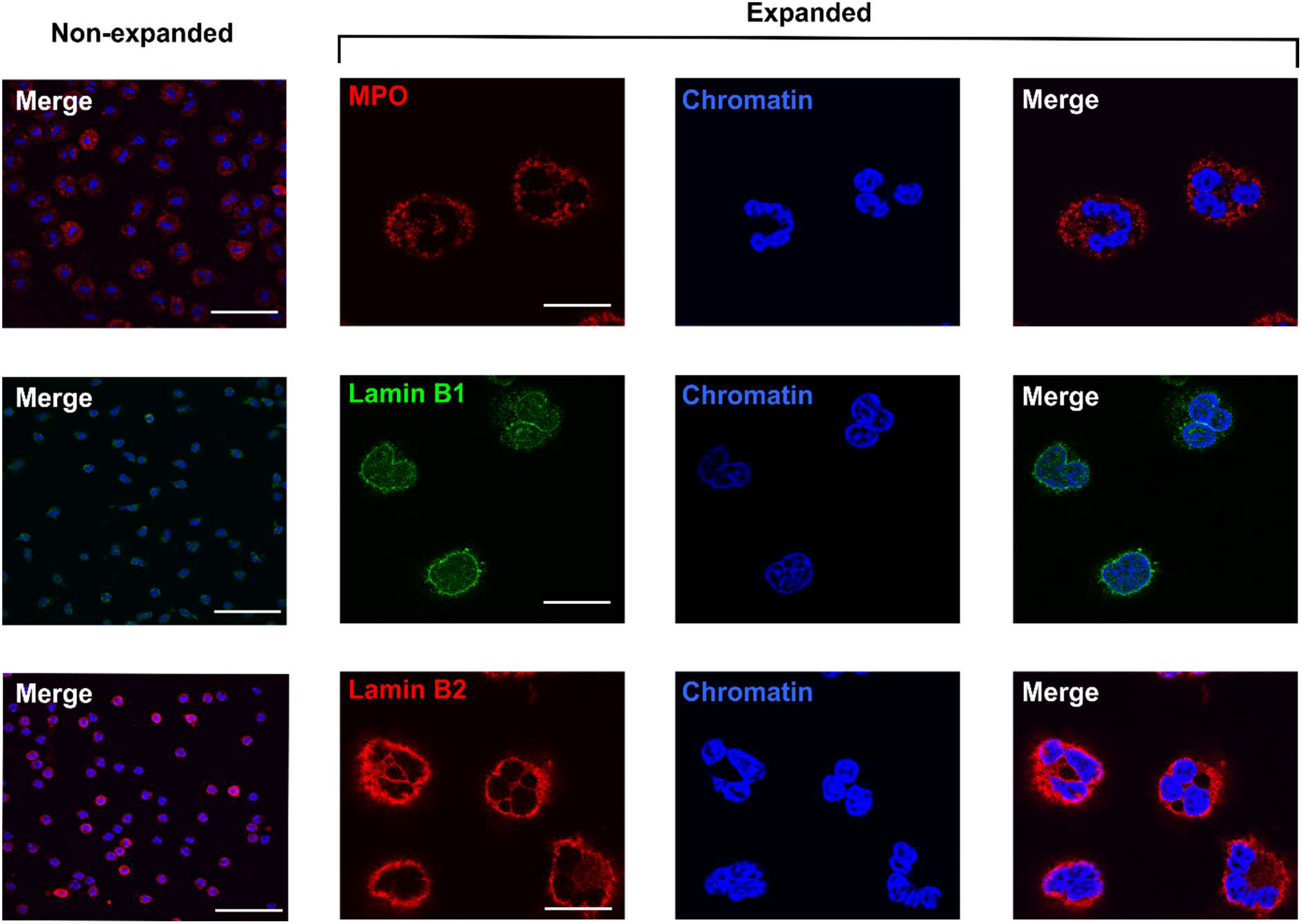
Expansion microscopy and immunofluorescence staining of neutrophil cytoplasmic and nuclear lamina proteins. MPO (first row) was stained as a characteristic cytoplasmic protein, and Lamin B1 (second row) and Lamin B2 (third row) as components of the nuclear lamina of neutrophils. Column 1 shows nonexpanded cells. Columns 2 through 4 show expanded cells, labelled as indicated. Scale bars = 50 μm.

Thus, expansion microscopy is well suited for staining a large range of epitopes inside neutrophils and can serve as general tool to study these cells. Similar to the case for histone H1 (figure 5) the increased amount of spatial information allowed us to learn more about the localization of certain proteins. NPM1 shows punctate agglomerations inside the nucleus that were not visible in the non-expanded cells. They could be attributed to NPM1-related liquid liquid phase separation (LLPS) and could play a role for gene regulation.

**Figure 7:**
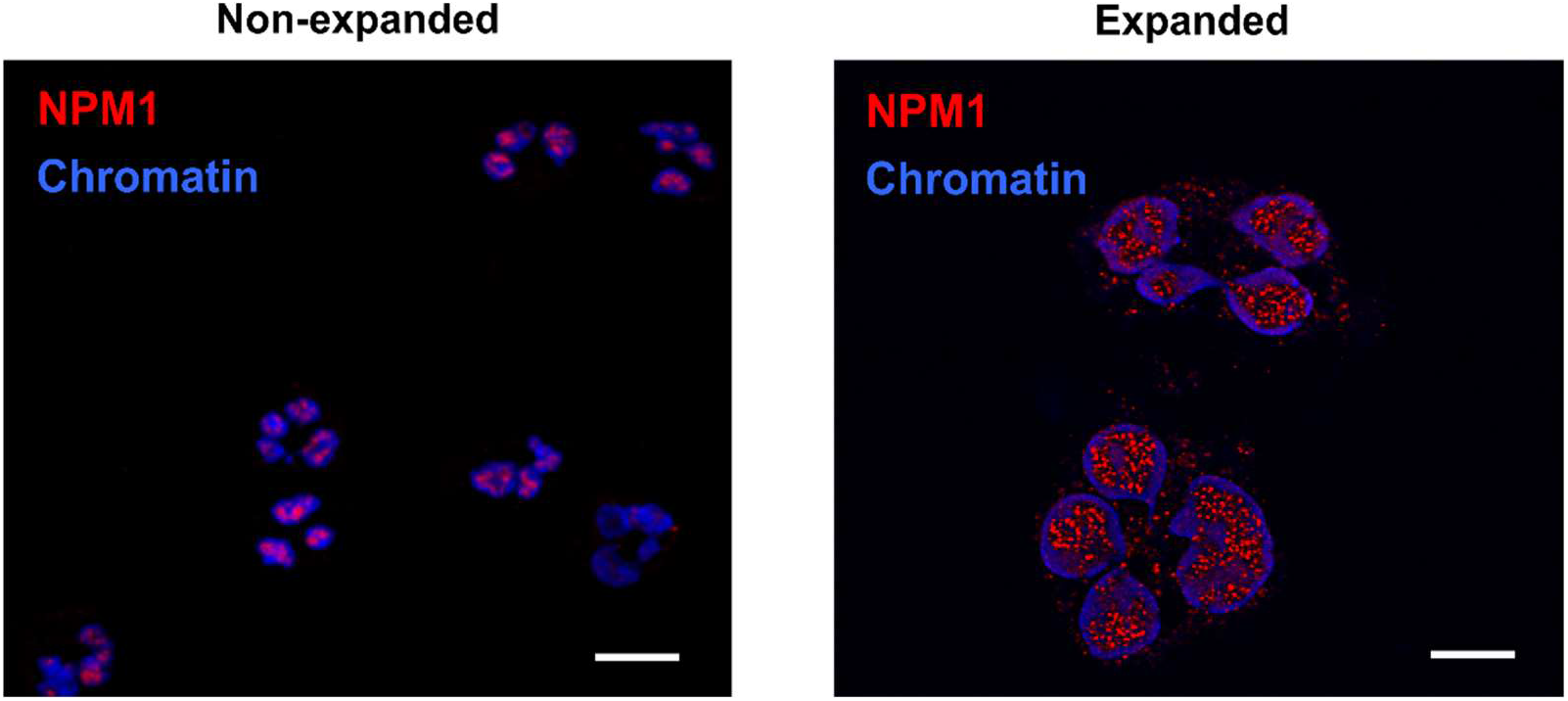
Expansion microscopy reveals punctate protein agglomerations in the neutrophil nucleus. NPM1 was stained as a nuclear protein involved in unique higher-order structural conformations. Left shows non-expanded cells and right shows expanded cells. The enhancement in resolution shows that NPM1 is not evenly distributed and appears as puncta within the chromatin of the nucleus. Scale bars = 10 μm.

## Discussion

Neutrophils are highly complex cells of the innate immune system. While originally considered a homogeneous population with a highly conserved repertoire of immune defense mechanisms, recent research has been shedding light on neutrophil heterogeneity and functional versatility. Indeed, neutrophils have emerged as key players not only during acute and chronic inflammation but also in malignant diseases and have thus been propelled into the focus of inflammation and cancer research. Of note, the neutrophil nucleus provides neutrophils with unique properties and abilities, including the comparatively high propensity for migration.

The rekindled interest in neutrophil biology has revealed the need for efficient methods to characterize neutrophil composition at high resolution. In principle, this can be accomplished with super-resolution techniques that are costly, time-consuming, and require specialized equipment. For this reason, we developed a novel, low-cost and generally accessible approach to super-resolution fluorescence images of neutrophils. Instead of relying on optical techniques to circumvent the limit of resolution, expansion microscopy enlarges the cellular structures by embedding the cells in a polyacrylamide hydrogel and then letting the hydrogel swell. Importantly, we have shown that the swelling process enlarges structures in a homogeneous manner without altering eccentricity (Figure 2). This is especially important in light of the lobulated morphology of nuclei in neutrophils. While the area is increased, the spatial relationship between structures is maintained. This was not only true for intact neutrophils but also for NETs, highlighting the general suitability of this method even for fragile structures.

To quantitate the improvement in resolution gained by expansion microscopy, we stained chromatin/DNA by a Hoechst dye and histone H1 or NPM1 by a fluorescently labeled antibody. As it was not possible to directly compare the same cell in an expanded versus a non-expanded state, we simulated a 4.9-fold decreased resolution in images of expanded cells by binning of 4 pixels (Figure 3 binned) to provide a direct measure for the improvement of resolution. Of note, this measure is similar to the gain in resolution.

We were able to show that the density of information across the neutrophil nucleus strongly increased by expansion microscopy (Figures 3A and 3B), as depicted in line scans along the axis of the nucleus. In the expanded cell, the line scan had shown a much higher amplitude and frequency of excursions as a measure for the gain of structural information. Expansion mproved the (theoretical) resolution from 163 nm to 34 nm (Hoechst dye).

Additionally, we calculated optical colocalization in non-expanded cells and expanded cells. In the non-expanded cells, differences in distribution cannot be assessed due to the limit in resolution, resulting in a Pearson’s coefficient closer to 1. Expansion microscopy significantly lowered the Pearson’s coefficient, as shown by Mann-Whitney U tests in Figure 4. Although chromatin and histone H1 or NPM1 are located together in the nucleus, their distribution is not identical (Figures 3A, 3B, and 7), which is in line with previously published literature^46,47^. These differences were suitably imaged by expansion microscopy, where conventional microscopy with non-expanded cells failed to show differences.

As a further proof of suitability of this method for neutrophils, we stained different neutrophil structures such as the nuclear lamins B1 and B2 and the cytoplasmic protein MPO (Figure 6), in addition to histone H1 and NPM1 (Figures 3, 5, and 7). All structures were imageable by the same expansion microscopy protocol. Strikingly, NPM1 appears as punctate ‘mininucleoli’ upon expansion, which has been discussed in the literature but not revealed in this detail^48^. Expanded histone H1 also strongly appeared in clearly defined border zones, often described as lamina-associated domains between heterochromatin and the nuclear lamina^49^. However, it should be noted that in the case of lamin B2 staining was somewhat blurred around the nucleus, especially in contrast to the very clear yet weaker lamin B1 staining. It is conceivable that the expansion process also led to a widening of the nuclear lamina and/or a certain washing out of the fluorophores. In the case of lamin B1, we observed the need for relatively high amounts of antibodies to achieve a good signal. As the expansion process does not increase epitopes but, conversely, “dilutes” them within the hydrogel, fluorophores with a good intensity are of high importance for this method. This is also reflected in the intensity of the staining in the line scans of Figure 3, as well as the lower signal intensities for most labels. Thus, expansion microscopy comes with the price of losing staining intensity while gaining resolution. In any case, the staining procedure should be adjusted for any new antibody/fluorophore that is to be used in combination with expansion microscopy, especially if one aims to study proteins that are not expressed in large quantities in the cell.

## Conclusion

In summary, we have demonstrated expansion microscopy for human neutrophils for different epitopes and fluorescent labeling strategies, including dyes (Hoechst) and antibody-based labeling. The increase in resolution allowed us to achieve a deeper understanding of the nuclear morphology and the nanoscale topography of chromatin and nuclear proteins. Therefore, expansion microscopy allows to better understand the complex and dynamic structure of nuclei of highly dynamic cells such as human neutrophils.

## Supporting material

Supporting material includes six figures: photographs of the gel handling technique (Supplementary Figure 1), photographs of the gel expansion time series (Supplementary Figure 2), diameter of granular particles of histone staining (Supplementary Figure 3), maximum intensity boxplots in a time series (Supplementary Figure 4), immunofluorescence staining of histone H1 as a nuclear border protein (Supplementary Figure 5), and isotype control images for the immunofluorescence stainings (Supplementary Figure 6).

## Acknowledgements

Funded by the Deutsche Forschungsgemeinschaft (DFG, German Research Foundation) under Germany’s Excellence Strategy-EXC 2033-390677874-RESOLV. We acknowledge support by the DFG via the Heisenberg program (S.K.). L.E. received support from the DFG (ER 321/2-1). J.S.H. is a member of the CiM-IMPRS graduate program Münster (CiM). This work is supported by the “Center for Solvation Science ZEMOS” funded by the German Federal Ministry of Education and Research BMBF and by the Ministry of Culture and Research of Nord Rhine-Westphalia.

## Author contributions

S.K. and L.E. designed the project; J.S.H., L.R., and T.H.B. performed research; J.S.H., L.S., L.R., and T.H.B. analyzed data; J.S.H., L.S., L.R., and E.N. generated figures; J.S.H., L.S., L.R., E.N., S.K., and L.E. wrote the manuscript.

