## Supplementary figures for "Expansion microscopy of nuclear structure and dynamics in neutrophils"

* Equal contribution

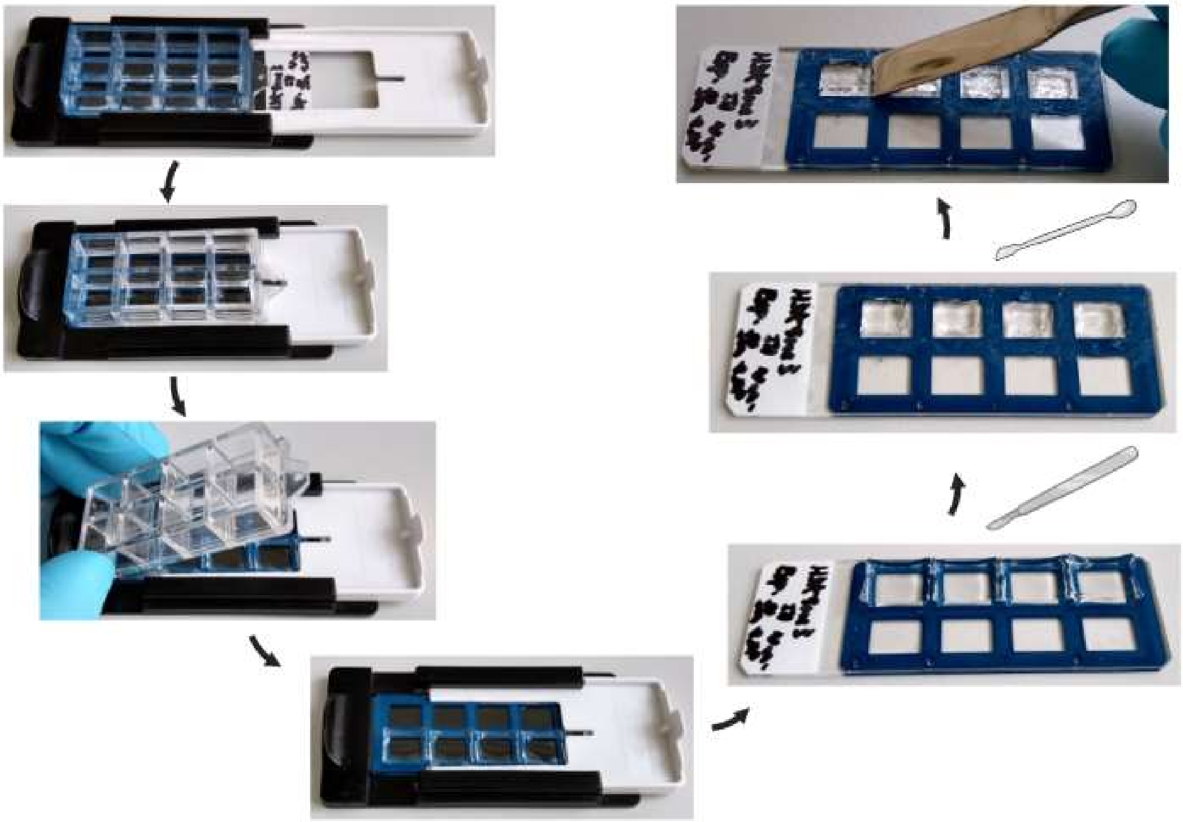

***Supplementary Figure 1: Photographs of expansion microscopy technique (8-well format):*** *Example images of gel processing and removal from 8-well chamber slide (Nunc Lab Tek II). Following chamber removal, gel edges were removed using a disposable scalpel and then gels removed from the slide using a flat spatula. Image created in BioRender.com.*

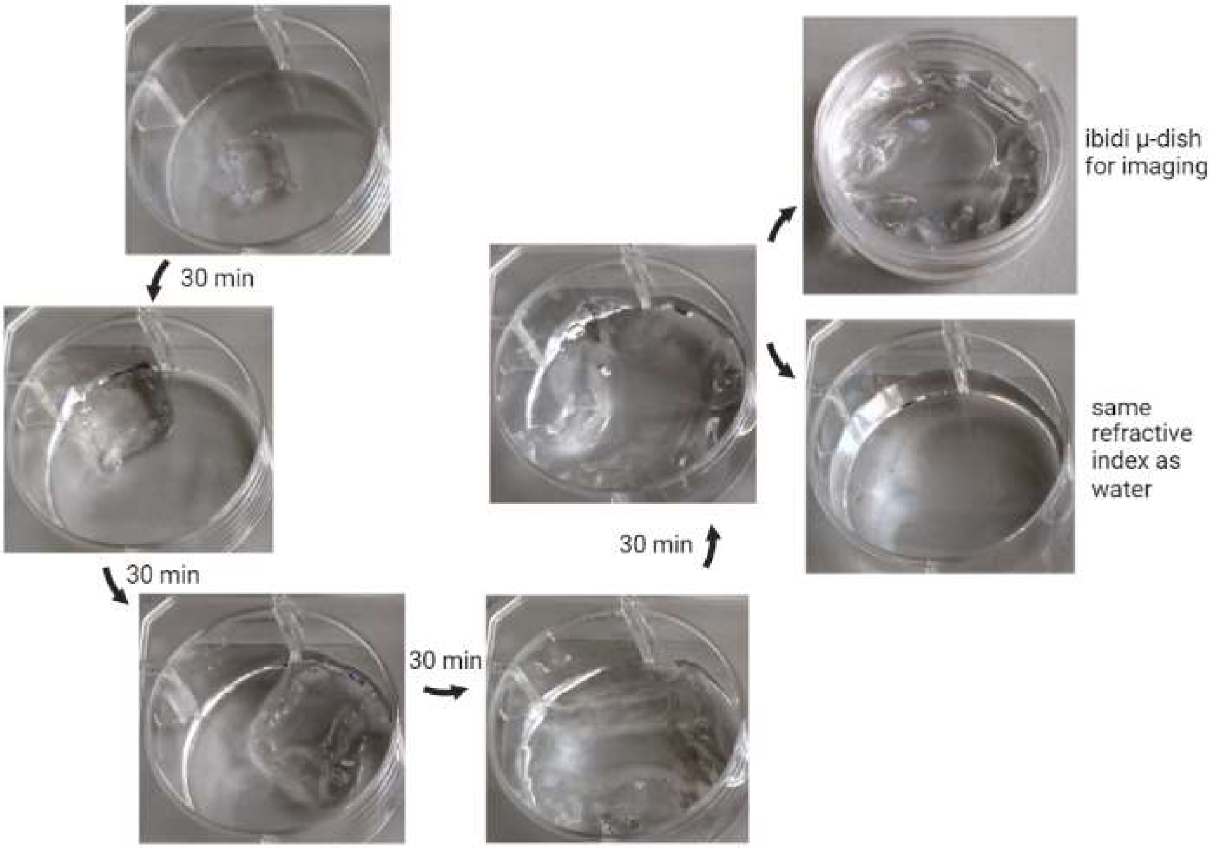


***Supplementary Figure 2: Photographs of expansion time series.*** *Image series showing expansion of gel following addition and exchange of distilled water (30 minutes between each, pictures taken during water exchanges). An ibidi µ-dish was used for imaging, facilitating simple gel handling with higher objectives. Final image shows expanded gel in water, demonstrating the gel having matched the refractive index of water. Image created in BioRender.com.*

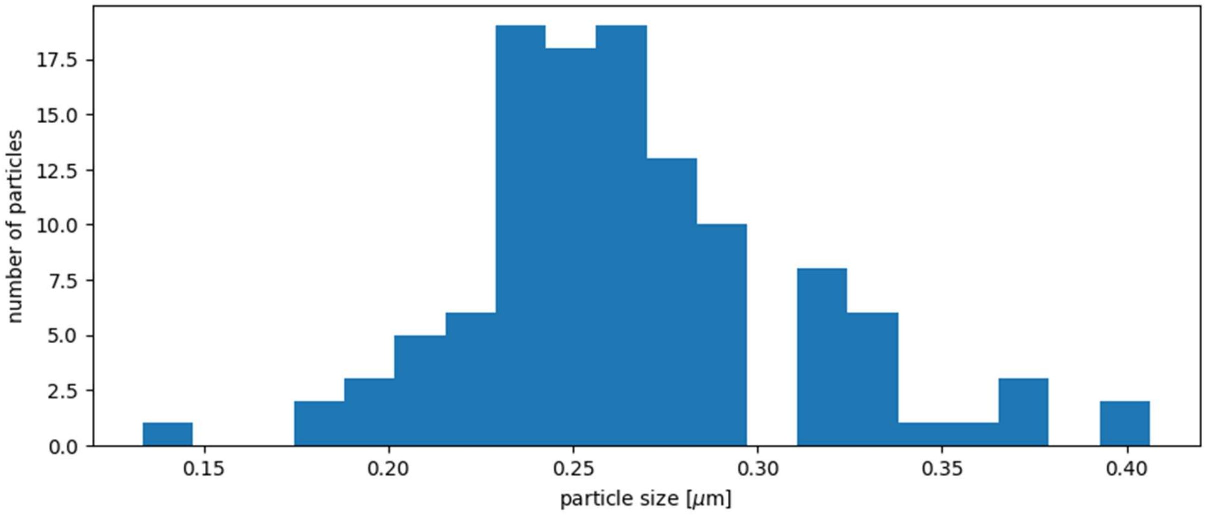


***Supplementary Figure 3: Histogram of granular particle diameters**: The granular particle diameter of the histone H1 staining of a neutrophil after 150min is shown. The mean diameter is 266nm ± 45nm which corresponds to 54.29 nm ± 9.18 nm before expansion. The small granular structures are without expansion microscopy below the resolution limit and would not be visible.*

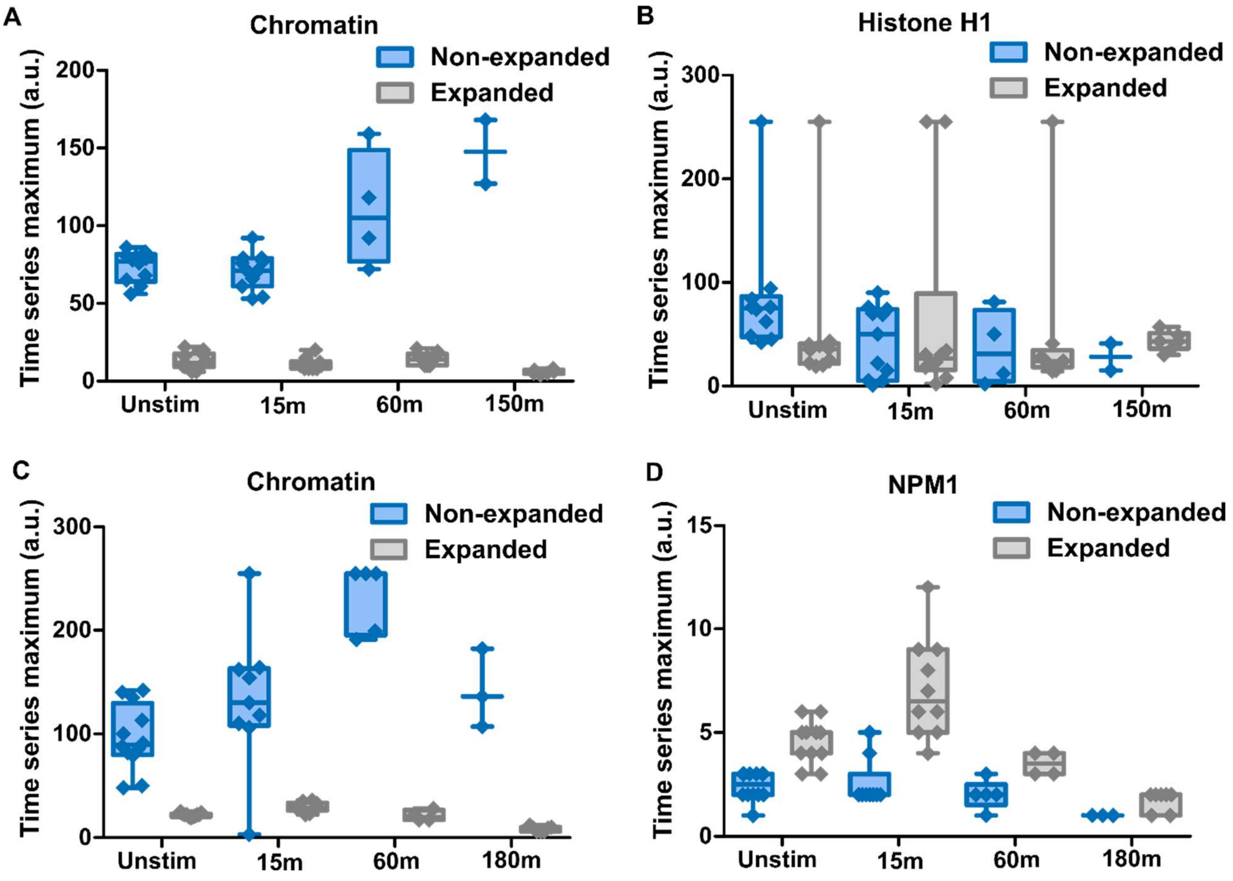


***Supplementary Figure 4: Maximum intensity of paired chromatin and histone H1 or NPM1 in a time series****: Boxplots of the maximum intensity for chromatin and histone H1 (**A**, **B**) and chromatin and NPM1 (**C**, **D**). Plots are paired and respectively labelled as non-expanded and expanded for each. Whiskers represent the maximum and minimum for each boxplot.*

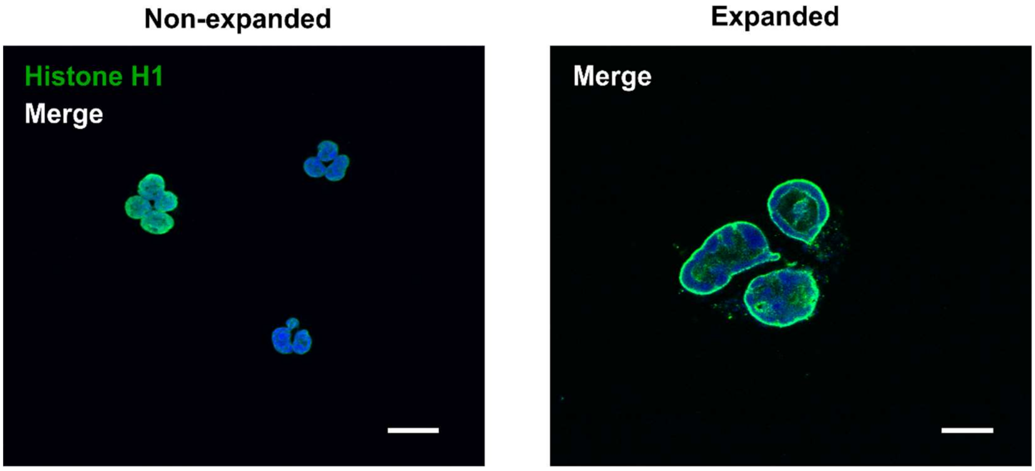


***Supplementary Figure 5: Expansion microscopy and immunofluorescence staining of neutrophil nuclear border proteins**. Histone H1 was stained as a nuclear protein involved in unique higher-order structural conformations. Left shows non-expanded cells and right shows expanded cells. Scale bars = 10 µm.*

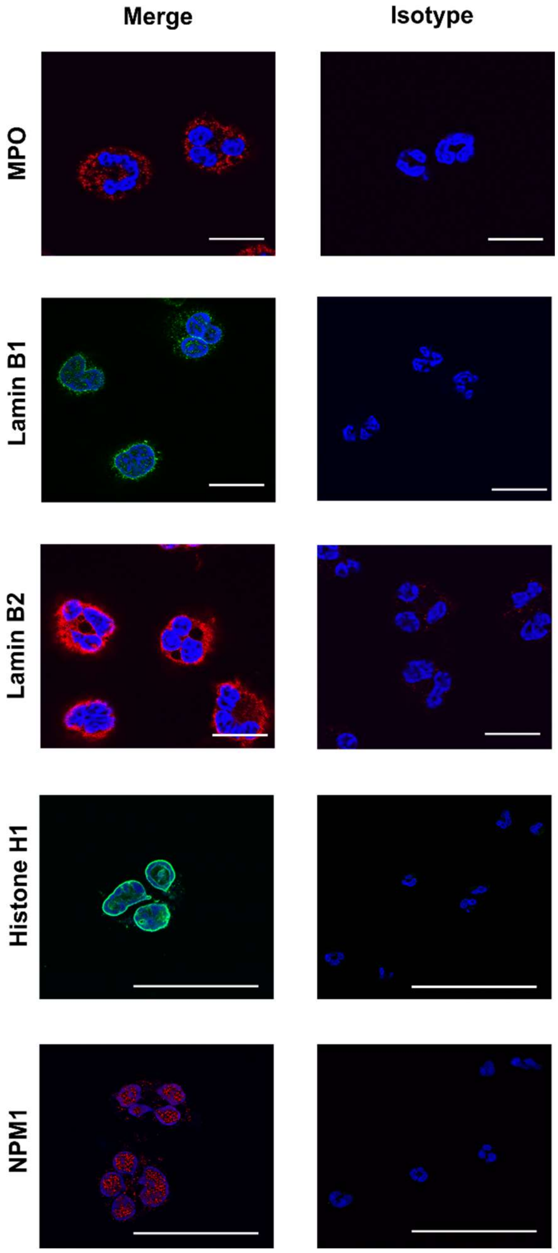


***Supplementary Figure 6: Immunofluorescence staining of neutrophil structures, including cytoplasmic and nuclear proteins****: Myeloperoxidase (MPO, first row) as a cytoplasmic protein, Lamin B1 (second row) and Lamin B2 (third row) as components of the nuclear lamina of neutrophils, and histone H1 (fourth row) and nucleophosmin (NPM1, fifth row) as nuclear proteins. Column 1 shows expanded chromatin and stained structures merged. Column 2 shows the respective isotype controls for these structures. Scale bars = 50 µm.*
